# Infection with CagA^+^ *Helicobacter pylori* induces epithelial to mesenchymal transition in human cholangiocytes

**DOI:** 10.1101/2020.04.28.066324

**Authors:** Prissadee Thanaphongdecha, Shannon E. Karinshak, Wannaporn Ittiprasert, Victoria H. Mann, Yaovalux Chamgramol, Chawalit Pairojkul, James G. Fox, Sutas Suttiprapa, Banchob Sripa, Paul J. Brindley

**Affiliations:** Department of Microbiology, Immunology and Tropical Medicine, and Research Center for Neglected Tropical Diseases of Poverty, School of Medicine & Health Sciences, The George Washington University, Washington DC, 20037, USA; Tropical Disease Research Laboratory, Faculty of Medicine, Khon Kaen University, Khon Kaen 40002, Thailand; Department of Pathology, Faculty of Medicine, Khon Kaen University, Khon Kaen, 40002, Thailand; Division of Comparative Medicine, Massachusetts Institute of Technology, Cambridge, 02139, MA

**Keywords:** *Helicobacter pylori*, cholangiocyte, epithelial-to-mesenchymal transition

## Abstract

Recent reports suggest that the East Asian liver fluke, *Opisthorchis viverrini*, infection with which is implicated in opisthorchiasis-associated cholangiocarcinoma, serves as a reservoir of *Helicobacter pylori*. The opisthorchiasis-affected cholangiocytes that line the intrahepatic biliary tract are considered to be the cell of origin of this malignancy. Here, we investigated interactions *in vitro* among human cholangiocytes, a CagA-positive strain of *Helicobacter pylori*, and the related bacillus, *Helicobacter bilis*. Exposure to increasing numbers of *H. pylori* at 0, 1, 10, 100 bacilli per cholangiocyte induced phenotypic changes including the profusion of thread-like filopodia and a loss of cell-cell contact, in a dose-dependent fashion. In parallel, following exposure to *H. pylori*, changes were evident in levels of mRNA expression of epithelial to mesenchymal transition (EMT)-encoding factors including snail, slug, vimentin, matrix metalloprotease, zinc finger E-box-binding homeobox, and the cancer stem cell marker CD44. Transcription levels encoding the cell adhesion marker CD24 decreased. Analysis to quantify cellular proliferation, migration and invasion in real time using the xCELLigence approach revealed that exposure to ≥10 *H. pylori* stimulated migration and invasion by the cholangiocytes through an extracellular matrix. In addition, 10 bacilli of CagA-positive *H. pylori* stimulated contact-independent colony establishment in soft agar. These findings support the hypothesis that infection with *H. pylori* contributes to the malignant transformation of the biliary epithelium.

## Introduction

There is increasing evidence that the East Asian liver fluke *Opisthorchis viverrini* may serve as a reservoir of *Helicobacter* which, in turn, implicates *Helicobacter* in pathogenesis of opisthorchiasis-associated cholangiocarcinoma (CCA) [1-5]. The International Agency for Research on Cancer of the World Health Organization classifies infection with the liver flukes *O. viverrini* and *Clonorchis sinensis* as well as *H. pylori* as Group 1 carcinogens [6]. In northern and northeastern Thailand and Laos, opisthorchiasis is the major documented risk factor for CCA [6-9]. Given the elevated prevalence of CCA in this region where infection with liver fluke prevails, and given evidence of linkages between infection by *Helicobacter* during opisthorchiasis, these two biological carcinogens together may orchestrate the pathogenesis of opisthorchiasis and bile duct cancer. Indeed, it has been hypothesized that the association of *Helicobacter* and its virulence factors, including during opisthorchiasis, underlies much biliary tract disease including CCA in liver fluke-endemic regions [10]. Infection with species of *Helicobacter* causes hepatobiliary disease that can resemble opisthorchiasis [5, 11-14]. Lesions ascribed to liver fluke infection including cholangitis, biliary hyperplasia and metaplasia, periductal fibrosis and CCA may be, in part, *Helicobacter*-associated hepatobiliary disease.

With respect to the gastric epithelium and the association with stomach adenocarcinoma, *H. pylori* colonizes the mucosal layer [6], adheres to the epithelium through bacterial adhesins with cellular receptors [15], from where its virulence factors stimulate a cascade of inflammatory signaling, anti-apoptosis, cell proliferation and transformation pathways by its virulence factors [16-18]. Fox and coworkers were the first to speculate that *H. pylori* also can cause hepatobiliary diseases in humans [19]. This is the dominant species among the genus *Helicobacter* detected in the context of hepatobiliary disease [20] and *H. pylori* is detected more frequently in cases with CCA or hepatocellular carcinoma (HCC) than in those with benign tumors and other control groups [21, 22], suggesting a positive correlation between *H. pylori* infection and hepatic carcinogenesis. In addition to the extensive literature on the interactions between *H. pylori* and gastric cells [17, 23], interactions between biliary epithelium and this bacillus have been reported. Among these, *in vitro* studies revealed that *H. pylori* induces multiple effects in CCA cell lines, including inflammation (IL-8 production), cell proliferation and apoptosis. Even at low multiplicity of infection, *H. pylori* induces pro-inflammatory cytokine and cell proliferative responses in CCA cell lines [4, 24], and even small numbers of bacilli that likely reach the biliary tract routinely may be sufficient to promote inflammation and transformation of the biliary epithelia [24].

Commensalism involving *H. pylori* and *O. viverrini* may have evolved and may facilitate conveyance of the bacillus into the biliary tract during the migration of the juvenile fluke following ingestion of the metacercaria in raw or undercooked cyprinid fish and establishment of liver fluke infection [1, 25]. Since curved rods resembling *Helicobacter* have been documented in the digestive tract of *O. viverrini* [1] and, given the low pH of the gut of the fluke, the *H. pylori* rods or spores might be transported from the stomach to the duodenum by the migrating larval parasite. The curved, helical *H. pylori* rod attaches to a cholangiocyte, which internalize in similar fashion to its colonization of mucous-producing cells of the gastric epithelia [17, 26, 27]. Transition of epithelial cells to mesenchymal cells during disease and epithelial to mesenchymal transition (EMT) during development and wound healing follow evolutionary conserved routes with well-characterized morphological and other phenotypic hallmarks [28]. These phenotypes are showcases during malignant transformation of the gastric mucosa resulting from infection with *H. pylori* [29]. Here, we investigated interactions between CagA^+^ *H. pylori*, the related species *H. bilis* [30], and human cholangiocytes. Infection with CagA^+^ *H. pylori* induced EMT in the H69 cell line of human cholangiocytes [31, 32].

## Materials and Methods

### Cell lines of human cholangiocytes

The immortalized intrahepatic cholangiocyte cell line, H69 [31, 32], Cellosaurus [33] identifier RRID:CVCL_812,1, and the primary cholangiocarcinoma cell line, CC-LP-1 [34, 35] Cellosaurus RRID CVCL_0205, were obtained as described [34-37]. In brief, H69 cells were cultured in Dulbecco’s Modified Eagle’s Medium (DMEM) (Thermo Fisher Scientific, Inc.), DMEM/F12 (Millipore-Sigma, St. Louis, MO) supplemented with 10% fetal bovine serum (FBS)(Invitrogen, Thermo Fisher Scientific, Inc.) and adenine, insulin, epinephrine, triiodothyronine/transferrin, hydrocortisone, epidermal growth factor and penicillin/streptomycin (all from Millipore-Sigma), as described [36]. CC-LP-1 cells were cultured in DMEM containing 10% FBS, L-glutamine and penicillin/streptomycin. Both cell lines were maintained at 37°C in humidified 5% CO_2_ in air. The cells were cultured to ∼80% confluence before co-culture with *H. pylori*. H69 cells at passages 10 to 22 only and CC-LP-1 cells at passages 5 to 10 were used in this study.

#### Helicobacter pylori and Helicobacter bilis

*Helicobacter pylori*, ATCC 43504, from human gastric antrum [38], and *H. bilis*, Fox *et al*. ATCC 49314, from intestines and liver of mice [39, 40], were obtained from the American Type Culture Collection (ATCC) (Manassas, VA) and maintained under a microaerobic atmosphere on trypticase soy agar with 5% sheep blood for 72 to 96 hours [41] (Becton Dickinson, Cockeysville, MD), incubated at 37°C with gentle agitation in a microaerobic atmosphere, established with the BBL Campy Pack Plus system (Becton Dickinson). After 96 hours, each species of *Helicobacter* was harvested by scraping the colonies from the agar plate, and resuspension of the scraping in DMEM/F12 medium until an optical density at 600 nm of 1. 0 was reached, which corresponds to ∼1×10^8^ colony forming units (cfu)/ml [42-44]. Viability of the bacteria was confirmed by visual inspection for bacterial movement.

### Morphological assessment of cholangiocytes

H69 cells were co-cultured with *H. pylori* at increasing multiplicities of infection (MOIs) of 0 (no bacilli), 10 and 100 for 24 hours in serum-free media. After 24 hours, the morphology of the H69 cells was documented and images captured using a digital camera fitted to an inverted microscope (Zeiss Axio Observer A1, Jena, Germany).

### Cell scattering and elongation

H69 cells were seeded at 5 × 10^5^ cells/well in 6-well culture plates (Greiner Bio-One, Monroe NC). At 24 hours, the culture medium was exchanged for serum- and hormone-free medium containing *H. pylori* at MOI of 0, 10, and 100, respectively, and maintained for a further 24 hours. At that interval, the appearance including scattering of the cells in the cultures was documented, as above. Cell scattering was quantified by counting the total number of isolated, single cells per field in 10 randomly selected images at 5x magnification [45]. To assess elongation of cells, images were documented of the cells in ∼20 randomly selected fields of view at ×20 magnification, with two to seven cells per field. The length to width ratio of the isolated cells was established using the ImageJ software [46].

### *In vitro* wound healing assay

Monitoring cell migration in a two-dimensional confluent monolayer may facilitate characterization the process of wound healing with respect with exposure to *H. pylori* [47]. A sheet migration approach was used to assay wound closure [36, 48, 49] following exposure of the cholangiocytes to *H. pylori*. H69 cells infected with *H. pylori* at MOI of 0, 10 and 100 were cultured overnight in 6-well plates to allow cell adherence. A linear scratch to wound the monolayer was inflicted with a sterile 20 µl pipette tip. The dimensions of wound were documented at 0 and at 26 hours, and the rate of wound closure quantified by measurement of the width of the wound in the experimental and the control (MOI of 0) groups of cells [36, 37].

### Real time quantitative PCR (RT-qPCR)

Total RNAs from H69 cells were extracted with RNAzol (Molecular Research Center, Inc., Cincinnati, OH) according to the manufacturer’s instructions. The RNA Quantity and quality of the RNAs were established by spectrophotometry (NanoDrop 1000, Thermo Fisher Scientific). Total RNAs (500 ng) were reversed-transcribed using the iScript cDNA Synthesis Kit (Bio-Rad, Hercules, CA). Analysis of expression of six EMT-associated genes -vimentin, snail, slug, ZEB1, JAM1 and MMP7, and two cancer stem cell markers, CD44 and CD24 - was undertaken by quantitative reverse transcription-polymerase chain reaction (qRT-PCR) of total RNAs using an ABI7300 thermal cycling system (ABI) and the SsoAdvance SYBR green mixture (Bio-Rad). Signals were normalized to expression levels for GAPDH. The relative fold-change was determined by the ΔΔCt method [50]. Three biological replicates of the assay were undertaken. Table S1 provides the oligonucleotide sequences of the primers in the RT-qPCRs. The design of these primers for the human EMT and stem cell markers was undertaken used Primer-BLAST, https://www.ncbi.nlm.nih.gov/tools/primer-blast/index.cgi [51], with the genome of *Helicobacter* specifically excluded during the blast search. Further, before the RT-qPCR analysis was undertaken, conventional PCR was carried out using cDNA from H69 cells as the template along with these primers. Products were sized using ethidium bromide-stained agarose gel electrophoresis, which confirmed that the presence of amplicons of predicted sizes (not shown).

### Assessment of cell proliferation

Cell proliferation of H69 was assessed using the xCELLigence real-time cell analyzer (RTCA) DP system (ACEA Biosciences, San Diego CA), as described [36, 52-54]. H69 cells were fasted for 4 to 6 hours in 1:20 serum diluted medium, 0.5% FBS final concentration, as described [55, 56], after which cells were co-cultured of viable bacilli of *H. pylori* for 120 min before starting the assay. H69 cells were subsequently harvested in 0.25% trypsin-EDTA and then washed with medium. Five thousand H69 cells were seeded on to each well of the E-plate in H69 medium and cultured for 24 hours. The culture medium was removed, and the cells were gently washed with 1×PBS. The PBS was replaced with serum diluted medium (above). The cellular growth was monitored in real time with readings collected at intervals of 20 minutes for ≥ 60 hours. For quantification, the cell index (CI) [53] was averaged from three independent measurements at each time point.

### Cell migration and invasion in response to *H. pylori*

To monitor the rate of cell migration and/or invasion in real time, we used the xCELLigence DP instrument (above) equipped with a CIM-plate 16 (Agilent ACEA Biosciences, San Diego CA), which is an electronic Boyden chamber consisting of a 16-well culture plate in which each well includes an upper chamber (UC) and a lower chamber (LC) separated by a microporous (pore diameter, 8 μm) membrane. In response to a chemoattractant, cells may migrate/invade from the UC towards the membrane and the LC. Migrating cells contact and adhere to microelectronic sensors on the underside of the membrane that separates the UC from the LC, leading to change in relative electrical impedance [53, 54, 57]. To investigate migration, H69 and CC-LP-1 cells were infected with *H. pylori* at MOIs of 0, 1, 10, 50 and 100 respectively for 120 min before the start of the assay, after which the cells were harvested following treatment for 3 min with 0.25% trypsin-EDTA and washed with medium. Subsequently, 30,000 H69 or CC-LP-1 cells were seeded into the UC of the CIM-plate in serum-free medium. Complete medium (including 10 % FBS) was added to the LC. Equilibration of the loaded CIM-plate was accomplished by incubation at 37°C for 60 min to allow cell attachment, after which monitoring for migration commenced in real time for a duration of ∼96 hours with readings recorded at 20 min intervals. For quantification, CI was averaged from three independent measurements at each time point [58].

To investigate invasion, H69 or CC-LP-1 cells were infected with *H. pylori* at either MOI of 0 or 10 at 120 minutes before commencing the t assay. The base of the UC of the CIM plate was coated with 20% solution of a basement membrane matrix (BD Matrigel, BD Biosciences, San Jose, CA) in serum-free medium. A total of 30,000 of H69 or CC-LP-1 cells were dispensed into the UC in serum-free medium. The LC was filled with H69 or CC-LP-1 medium, as appropriate, containing 10% FBS. The loaded CIM-plate was sealed, inserted into the RTCA DP xCELLigence instrument at 37° C in 5% CO2 and held for 60 min to facilitate attachment of the cells, after which electronic recording was commenced. Monitoring for invasion in real time continued for ∼96 hours with the impedance value recorded at intervals of 20 min continuously during the assay. CI was quantified as above.

### Colony formation in soft agar

For the soft agar assay, H69 cells were infected with *H. pylori* or *H. bilis* at MOI of 0, 10, 50 and 100 at the start of the assay. H69 cells at 10,000 cells per well were mixed with 0.3% low melting temperature agarose (NuSieve GTG) (Lonza, Walkersville, MD) in complete medium, and plated on top of a solidified layer of 0.6% agarose in growth medium, in wells of 6- or 12-well culture plates (Greiner Bio-One). Fresh medium was added every three or four days for 28 days or until the colonies had established, at which time the number of colonies of ≥ 50 µm diameter was documented using a camera fitted to an inverted microscope (above) and quantified [59].

## Statistical analysis

Differences in cell elongation, cell scattering, wound healing and colony formation among experimental and control groups were analysed by one-way analysis of variance (ANOVA). Fold differences in mRNA levels, and real time cell migration and invasion through extracellular matrix were analysed by two-way ANOVA followed by Dunnett’s test for multiple comparisons. Three or more replicates of each assay were carried out. Analysis was accomplished using GraphPad Prism v7 (GraphPad, San Diego, CA). A *P* value of ≤ 0.05 was considered to be statistically significant.

## Results

### *Helicobacter pylori* induces epithelial to mesenchymal transition in cholangiocytes

H69 were directly exposed to increasing number of *H. pylori* at 0, 10 and 100 bacilli per cholangiocyte in 6-well plates. At 24 hours after addition of the bacilli, the normal epithelial cell appearance in tissue culture had altered to an elongated phenotype characterized by terminal thread-like filopodia and diminished cell-to-cell contacts (Figure 1A-C). The extent of the transformation was dose-dependent and, also, it indicated increased cell motility. The morphological change was quantified by measurement of the length to width ratio of the exposed cells. In a dose dependent fashion, the ratio increased significantly at both MOI of 10 and of 100 when compared with MOI of 0, *P* ≤ 0.01 (Figure 1D-F). In addition, the numbers of isolated, single cells increased significantly to the increasing length to width ratio of the *H. pylori*-exposed cholangiocytes, *P* ≤ 0.05 and ≤ 0.001 at MOI of 10 and 100, respectively (Figure 1G). The cholangiocytes displayed had a hummingbird phenotype-like appearance, reminiscent of AGS gastric line cells [29]. Moreover, compared to H69 cells cultured in medium supplemented with TGF-β at 5 ng/ml displayed cell aggregation and nodule formation (Figure S1).

**Figure 1.**
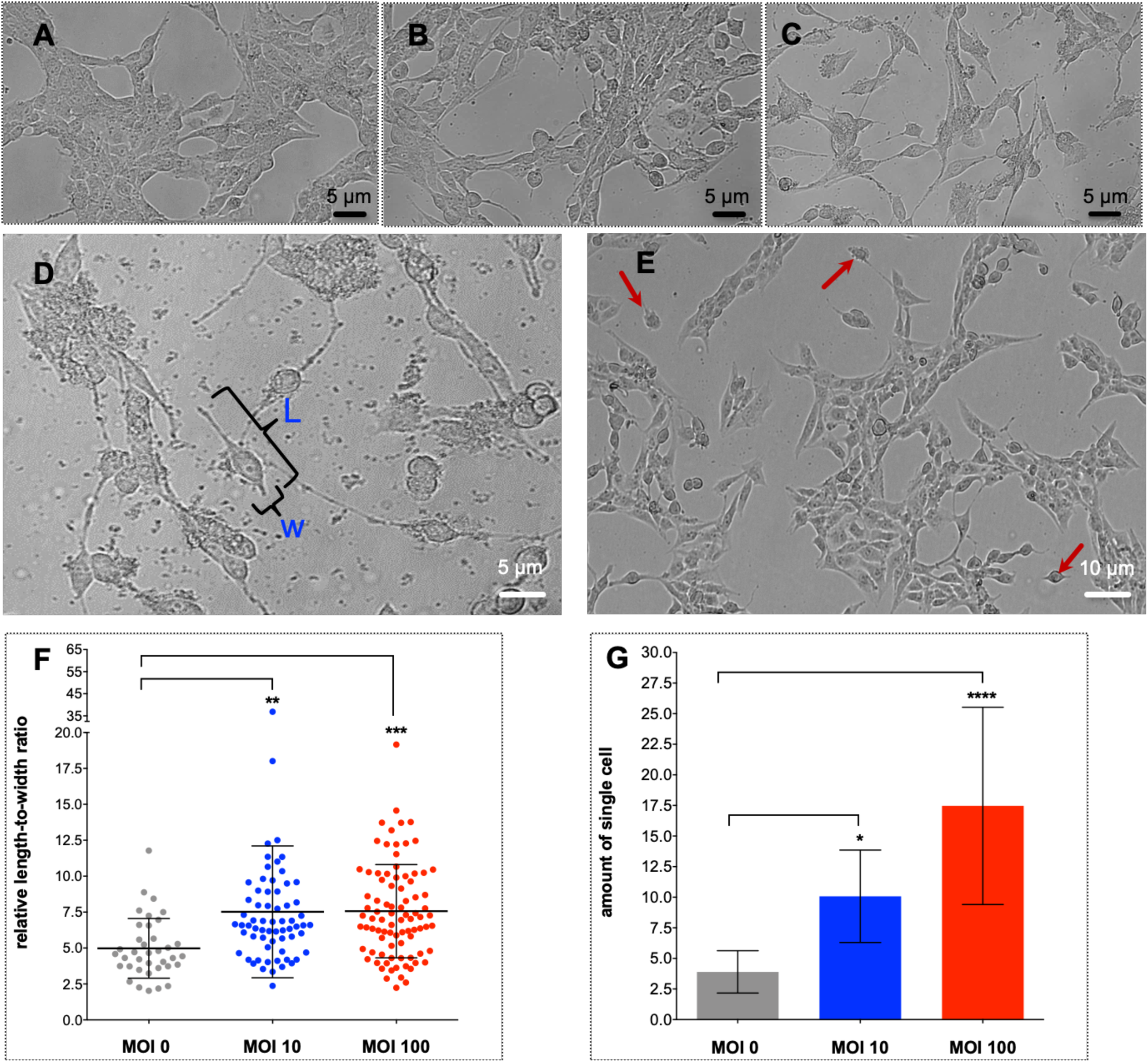
Exposure to CagA^+^ *Helicobacter pylori* ATCC 43504 induced morphological alteration in cholangiocytes including cell elongation, indicative of epidermal to mesenchymal transition. Panels A-C: photomicrographs documenting cell morphology of H69 cells exposed to increasing numbers of *H. pylori* at 0, 10, and 100 bacilli per cholangiocyte, respectively (left to right). The appearance of cells changed from an epithelial to a mesenchymal phenotype as evidenced by the loss of cell-cell contact, an elongated and spindle-shaped morphology, along with growth as individual cells by 24 hours following *H. pylori* infection in a dose-dependent manner. Scale bars, 5 μm (right), 20× magnification. The length-to-width ratio of single, isolated single cells were documented to determine cellular elongation and scattering (D, E). The number of elongated cells increased in a dose-dependent fashion in response to infection by *H. pylori* (F). By contrast, the number of isolated individual H69 cells, indicative of cell scattering, was also significantly increased in dose-dependent fashion (G). Data are presented as the mean ± standard error of three biological replicates. Means were compared using one-way ANOVA. Asterisks indicate levels of statistical significance of experimental compared to control groups at 24 hours; *, *P* ≤ 0.05; **, *P* ≤ 0.01; ***, *P* ≤ 0.001; ****, *P* ≤ 0.0001.

### EMT-associated-factors induced by exposure to *H. pylori*

Transcriptional dynamics of the apparent EMT was investigated using qRT-PCR of total cellular RNAs for six EMT-related factors including the mesenchymal marker vimentin, the transcription factors, Snail, Slug and ZEB1, the adhesion molecule JAM1 and the proteolytic enzyme, MMP7 [60-62]. The cancer stem cell markers, CD24 and CD44 [63] also were monitored. Levels of the fold-difference in transcription for each of the six markers increased in a dose-dependent fashion. The regulatory transcriptional factor, Snail was markedly up-regulated with highest level of fold difference, 88-, 656- and 5,309-fold at MOI of 10, 50 and 100, respectively, followed by MMP7, with 15-, 21-, and 44-fold at MOI of 10, 50 and 100, respectively. Transcription of Slug, ZEB1, vimentin and JAM1 also was up-regulated during bacterial infection. CD24 expression was down-regulated whereas CD44 was up-regulated, revealing a CD44^+^ high/CD24^+^ low phenotype in cells exposed to *H. pylori* (Figure 2; *P* ≤ 0.05 to ≤ 0.0001 for individual EMT markers and/or MOI level, as shown). A pattern of CD44^+^ high/CD24^+^ low expression is a cardinal character of cancer stem cell activity in gastric adenocarcinoma [63].

**Figure 2.**
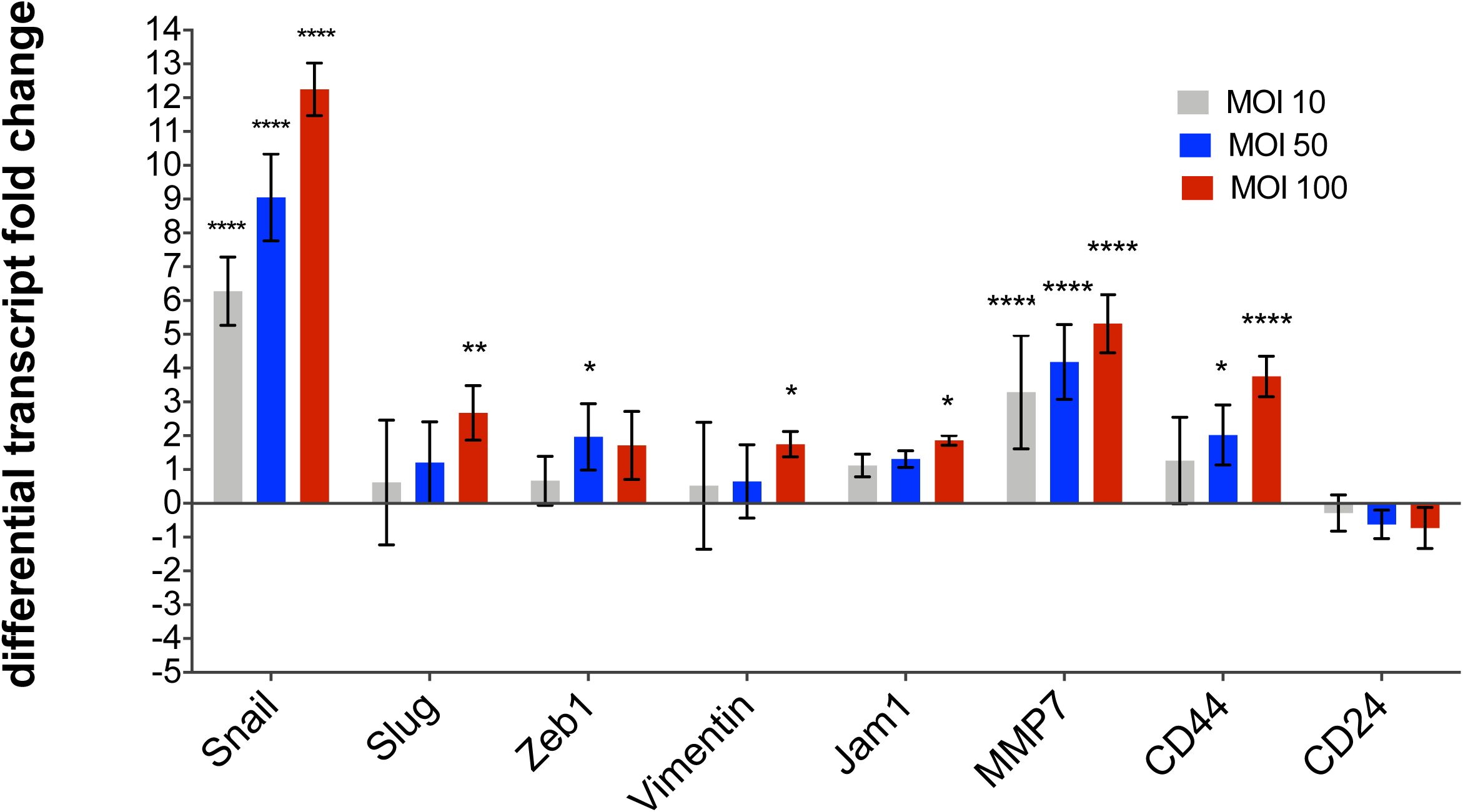
Differential transcript fold change of EMT-related and cancer stem cell marker genes after exposure to *Helicobacter pylori*. Messenger RNA expression of six EMT-related genes and two cancer stem cell markers were determined after 24 hours of infection. Expression of Snail, Slug, vimentin, JAM1, MMP7, and CD44 increased in a dose fashion whereas CD24 transcription decreased. Expression of the regulatory transcriptional factor, Snail was notably up-regulated by 6.27±1.02-, 9.05±1.28- and 12.25±0.78-fold at MOI 10, 50, and 100, respectively. MMP7 expression was markedly up-regulated by 3.3 to 5.3 fold at each MOI. Expression of each of Slug, ZEB1, vimentin, and JAM1 was up-regulated in a dose dependent fashion. Transcription of the cancer stem cell marker CD44 was significantly up-regulated by 2.02±0.88 fold at MOI 50 and by 3.75±0.60 fold at MO of 100 whereas no significant change was seen with CD24. Three biological replicates were carried out. The qPCR findings were normalized to the expression levels of GAPDH in each sample, with the mean ± S.D. values shown for the seven genes at each of MOI of 10, 50 and 100, and compared using a two-way ANOVA multiple comparison with a 95% confidence interval of difference.

### Exposure of cholangiocytes to *H. pylori* induces cellular migration, invasion and wound closure

Analysis in real time with a Boyden chamber-type apparatus revealed that exposure to *H. pylori* at MOI of 10 to 50 *H. pylori* significantly stimulated migration of H69 cells from 24 to 96 hours after starting the assay (Figure 3A; *P* values at representative times and MOIs). The effect was not evident at MOI 100. A similar assay was undertaken using the CC-LP-1 cholangiocarcinoma cell line but with at lower MOI. At MOI of 1, 10 and 50, *H. pylori* stimulated significantly more migration of CC-LP-1 cells from 20 to 40 hours after starting the assay (Figure 3B). Concerning invasion of extracellular matrix, both the CC-LP-1 and H69 cells migrated and invaded the Matrigel layer in the UC of the CIM plate at significantly higher rates than did the control cells, with significant differences evident from 48 to 96 hours (Figure 3, C,D). In addition, scratch assays revealed two-dimensional migration of H69 cells over 24 hours. Wound closure by H69 cells increased significantly to 19.47% at MOI of 10 (*P* ≤ 0.05) although an effect at MOI 100 was not apparent in comparison with the control group (Figure 4).

**Figure 3.**
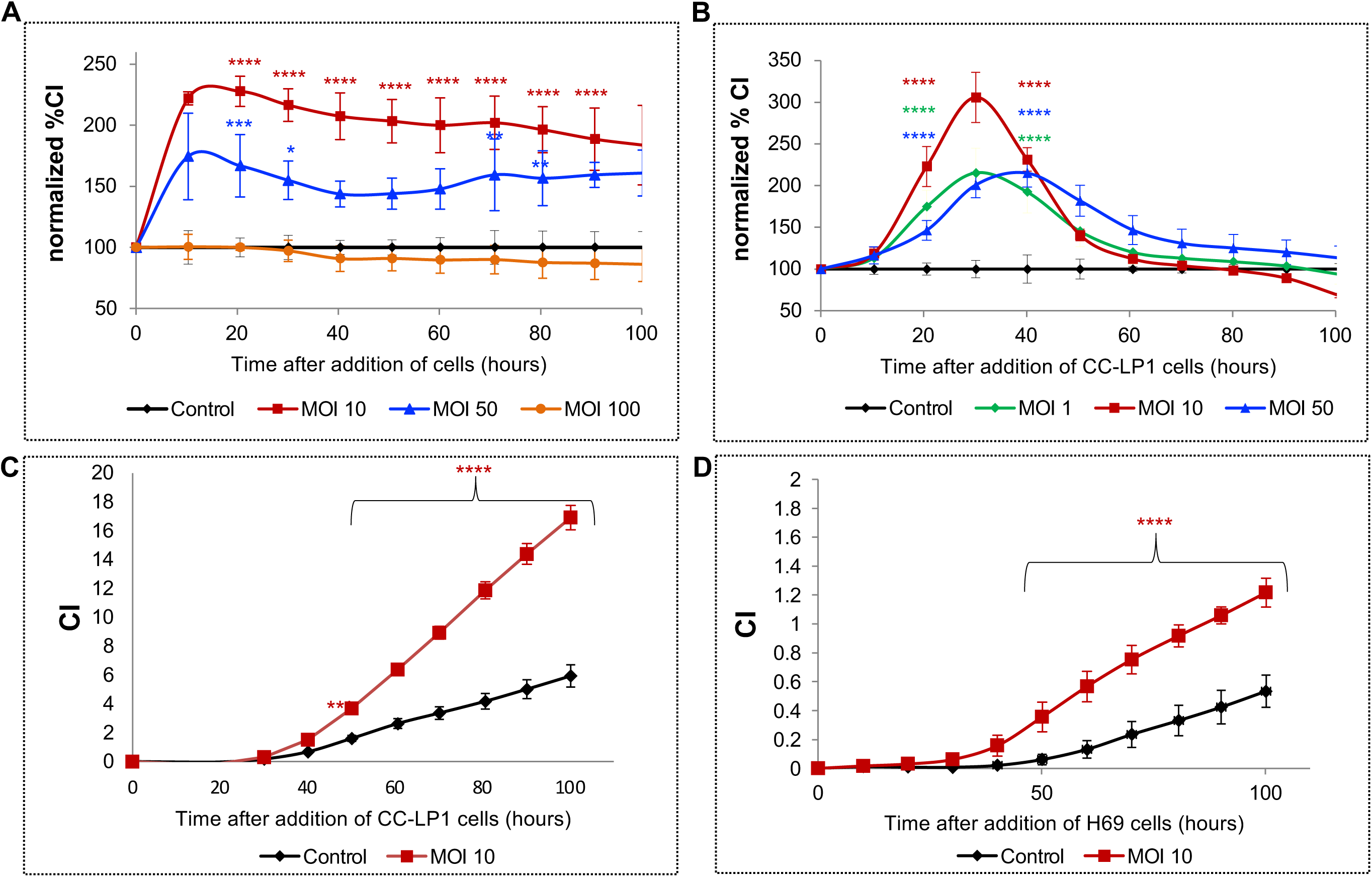
Exposing cholangiocytes to *H. pylori* induced migration and invasion through extracellular matrix. H69 cholangiocytes infected with *H. pylori* migrated through Matrigel, as monitored in real time over 100 hours using CIM plates fitted to a xCELLigence DP system. Cell migration and invasion were monitored for 100 hours with chemo-attractant in the lower chamber of CIM plate. H69 cells infected with MOI 10 exhibited the highest cell migration rate followed by MOI 50 whereas MOI 100 attenuated cell migration to a level comparable with non-infected control (A). Similarly, the cholangiocarcinoma cell line CC-LP-1 showed the highest cell migration rate when stimulated with MOI 10 followed by MOI 50. Moreover, CC-LP-1 cells significantly migrated faster even with at MOI of 1 (B). Invasion of the Matrigel extracellular matrix compared between MOI 10 and non-infected cells; invasion rates for CC-LP-1 and H69 cells significantly increased following exposure to *H. pylori* (C,D).

**Figure 4.**
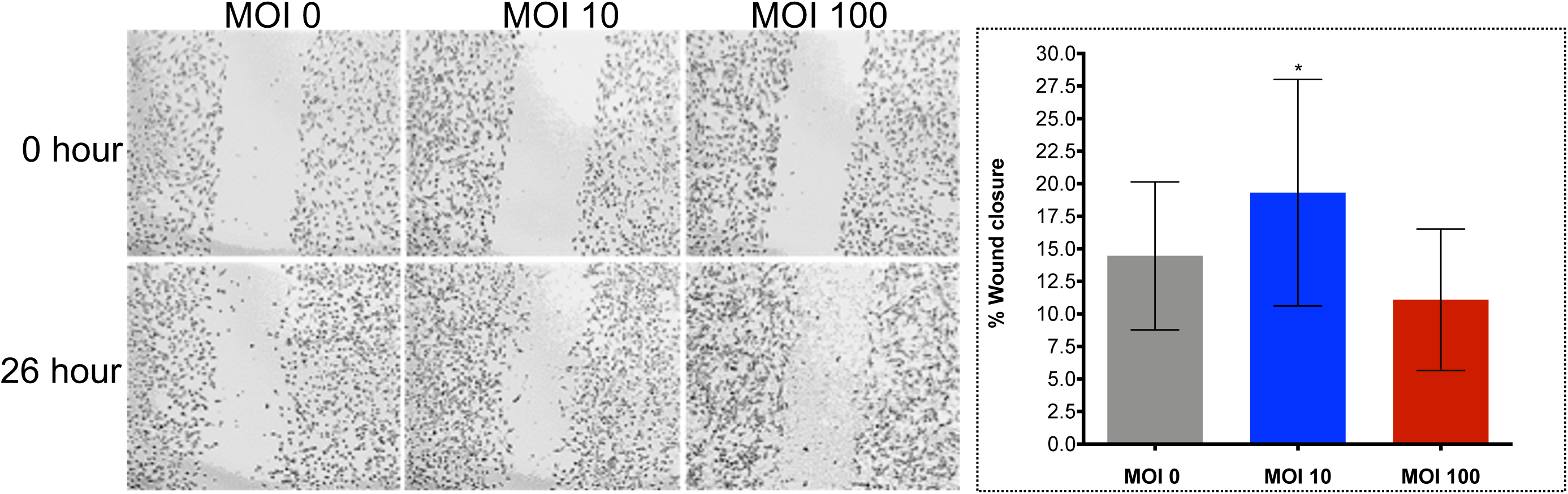
Wound healing in a two-dimensional *in vitro* assay revealed cell migration of H69 cholangiocytes infected with *H. pylori*. Wound closure was significantly increased to 19.47% at MOI 10 of *H. pylori* (*P* ≤ 0.05) by one-way ANOVA. An increase in wound closure was not apparent at MOI 100; 11.09% vs control, 14.47%. Micrographs at 0 (left) and 26 (right) hours; 5× magnification.

### *H. pylori* induces anchorage-independent colony formation

As a prospective biomarker of malignant transformation of the *Helicobacter*-infected cholangiocytes, anchorage independent colony formation by the cells was determined in a medium of soft agar. Following maintenance of the cultures for up to 28 days, counts of the numbers of colonies was revealed that a MOI of 10 *H. pylori* had induced a significant increase colony numbers of the H69 cells (Figure 5A, B). Significant differences from the control group were not seen at MOI 50 or 100. The size of colonies also increased in all groups exposed to *H. pylori* although this was statistically significant only at the MOI of 50 (Figure S2; *P* ≤ 0.05). By contrast to *H. pylori*, exposure of H69 cells to *H. bilis*, a microbe that naturally resides in the biliary tract and intestines [40, 64], decreased the number of colony formation in soft agar (Figure 5C). The neutral or even inhibitory effect of *H. bilis* on H69 was mirrored by the lack of cellular proliferation by H69 cells when monitored in real-time over 72 hours days at MOI of 0, 10, 50, and 100 *H. bilis* (Figure S3).

**Figure 5.**
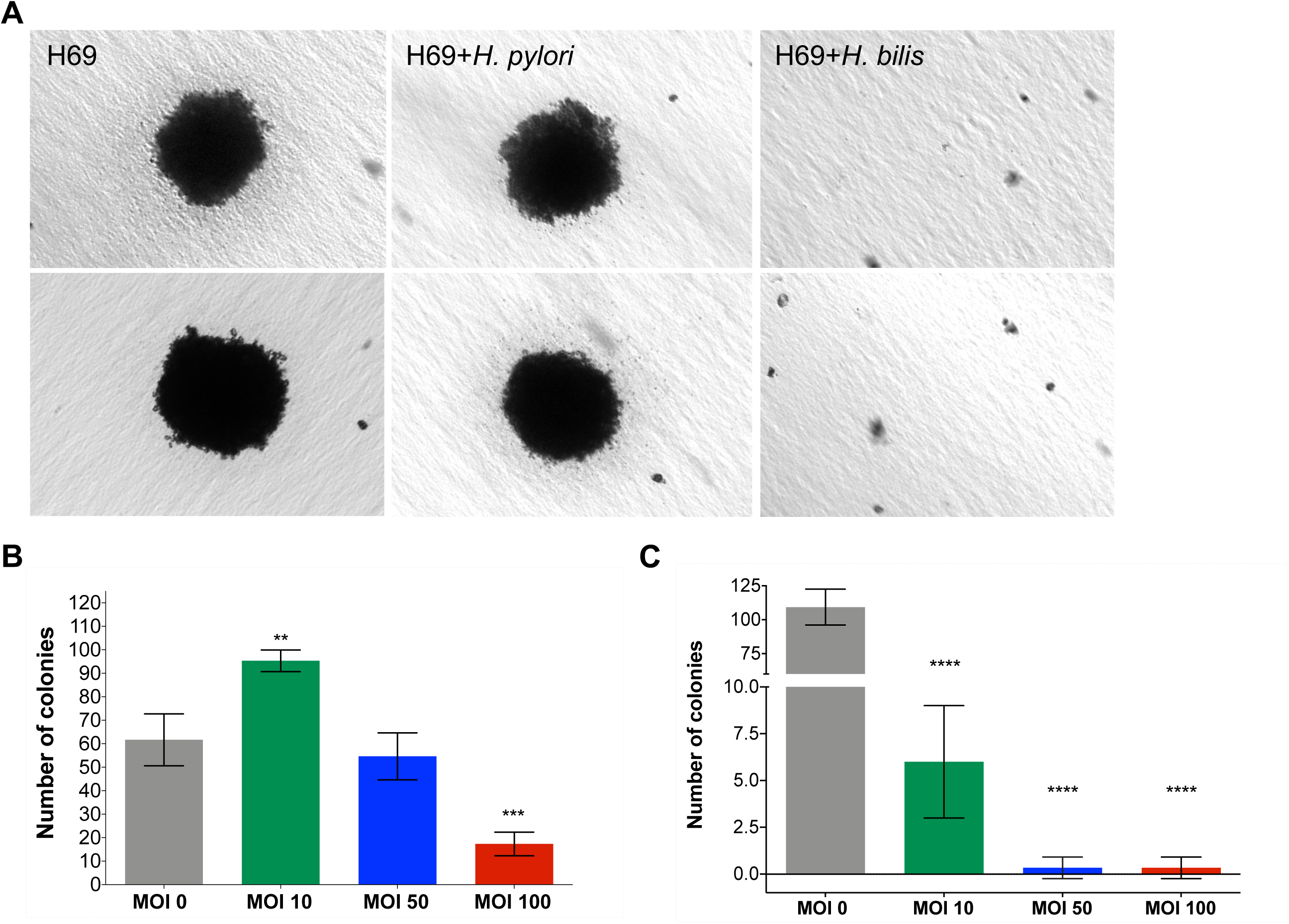
Anchorage-independent cell growth in soft agar revealed cellular transformation of cholangiocytes by *H. pylori*. Representative micrographs revealing the appearance of colonies of H69 cholangiocytes at 30 days following exposure to *H. pylori* and to *H. bilis*, as indicated (A). At MOI of 10, the number of H69 cell colonies increased significantly (*P* ≤ 0.01), whereas not at MOI of 50; by contrast, there was a significant decrease in colony numbers at MOI of 100, when compared with the non-infected control cells (B). Exposure of H69 cells to *H. bilis* resulted in markedly reduced numbers of colonies, in a dose-dependent fashion (C; *P* ≤ 0.001), indicating an inhibitory effect of *H. bilis* toward anchorage-independent cell growth and/or cellular transformation of human cholangiocytes (C). Three biological replicates were performed.

## Discussion

Numerous species of *Helicobacter* have been described [65] while *H. pylori* was the first prokaryote confirmed to cause gastric disease, including peptic ulcer, gastric mucosa-associated tissue lymphoma, and adenocarcinoma [38, 66-69]. Bacilli of *H. pylori* occur in the stomach in at least half of the human population. Transmission is from mother to child and by other routes.

The human-*H. pylori* association may be beneficial in early life, including contributions to a healthy microbiome and reduced early-onset asthma [70, 71]. Strains of *H. pylori* have been characterized at the molecular level, and classified as either CagA (cytotoxin-associated gene A)-positive or -negative. CagA, the major virulence factor of *H. pylori*, is a ∼140 kDa protein encoded on the Cag pathogenicity island, PAI. Cag PAI also encodes a type 4 secretion channel through which the virulence factor is introduced into the host cell [29]. CagA locates to the cell membrane, where it is phosphorylated by the Src kinases. Downstream of these modifications, the activated CagA orchestrates changes in morphology of the epithelial cell, which loses cell polarity and becomes motile, transforming in appearance to the hallmark ‘hummingbird-like’ phenotype [72-74].

The CagA oncoprotein is noted for its variation at the SHP2 binding site and, based on the sequence variation, is subclassified into two main types, East-Asian and Western CagA. East-Asian CagA shows stronger SHP2 binding and greater biological activity than the Western genotype. In East Asia, the circulation of *H. pylori* strains encoding active forms of CagA may underlie the elevated incidence of gastric adenocarcinoma [29]. In addition to the well-known association with gastric cancer, *H. pylori* has been associated with hepatobiliary disease [4, 5, 75]. Related species of *Helicobacter, H. hepaticus* and *H. bilis*, also associate with hepatobiliary disease [75-77]. Infection with the fish-borne liver flukes, *Opisthorchis viverrini* and *Clonorchis sinensis*, as well as the infection with *H. pylori* are all classified as Group 1 carcinogens by the International Agency for Research on Cancer [6]. Opisthorchiasis is a major risk factor for cholangiocarcinoma in northeastern Thailand and Laos [6-9]. In addition to gastric disease, infection with species of *Helicobacter* causes hepatobiliary tract diseases that can resemble opisthorchiasis [5, 8, 9, 11, 12]. Liver fluke infection can induce lesions in biliary system including cholangitis, biliary hyperplasia and metaplasia, periductal fibrosis and CCA. These lesions derive not only from liver fluke infection but perhaps are in part the consequence of hepatobiliary tract infection with *H. pylori. Helicobacter* may transit from the stomach to the duodenum and enter the biliary tree through the duodenal papilla and ampulla of Vater [26, 78], and indeed may be vectored there by the liver fluke, *O. viverrini* [1-3]. *H. pylori*-specific DNA sequences have been detected in CCA tumors and also from lesions diagnosed as cholecystitis/cholelithiasis in regions endemic for opisthorchiasis [5, 11]. Furthermore, serological findings have implicated infection with *H. pylori* as a risk for CCA in Thailand [12].

Here, we investigated interactions among a human cholangiocyte cell line, H69, a CCA cell line CC-LP-1, CagA^+^ *H. pylori*, and *H. bilis* bacilli. Infection of H69 with CagA^+^ *H. pylori* induced EMT in dose-dependent manner, which was characterized by cell elongation and scattering which, in turn, implicate increasing change in cell motility. This visible appearance of these infected H69 cells resembled the hummingbird phenotype of gastric epithelial cells after exposure *H. pylori* [28, 29, 79]. In the AGS cell line, delivery of CagA by the type IV secretion mechanism from *H. pylori* subverts the normal signaling leading to actin-dependent morphological rearrangement. The hitherto uniform polygonal shape becomes markedly elongated with terminal needle-like projections [80].

AGS cells infected with *H. pylori* demonstrating the hummingbird phenotype also display early transcriptional changes that reflect EMT [28]. H69 cells infected with *H. pylori* exhibited upregulation of expression of Snail, Slug, vimentin, JAM1, and MMP7 in a dose-dependent fashion, changes that strongly supported the EMT of this informative cholangiocyte cell line.

Likewise, Snail, an *E*-cadherin repressor, was markedly up-regulated, which indicated that this factor may be a key driver of EMT in cholangiocytes. CD44 expression was up-regulated in dose-dependent fashion, whereas CD24 decreased, showing a CD44^+^/CD24^-/low^ phenotype during infection. *H. pylori* may not only induce EMT but also contribute to stemness and malignant transformation of the cholangiocyte [63, 81].

Realtime cell monitoring revealed that the H69 and CC-LP-1 cells migrated when exposed to *H. pylori*. Moreover, 10 bacilli of *H. pylori* per biliary cell stimulated cellular migration and invasion through a basement membrane matrix, a behavioral phenotype also characteristic of EMT. In addition, following exposure to *H. pylori*, H69 cells responded with an anchorage-independent cell growth in soft agar, a phenotype indicative of the escape from anoikis and characteristic of metastasis [82]. The numbers of colonies of H69 cells in soft agar significantly increased following exposure to *H. pylori* at MOI of 10. By contrast, monitoring contact-independent cell growth in soft agar and also growth responses by H69 cells as monitored and quantified using the RTCA approach, both the numbers of colonies of H69 in soft agar and cell growth during the RTCA assay both decreased. The responses indicated that CagA^+^ *H. pylori*, but not *H. bilis*, displays the potential to induce neoplastic changes in cholangiocytes in like fashion to its carcinogenicity for the gastric epithelia.

The findings presented here notwithstanding, the report has some limitations. We utilized the ATCC 43504 CagA positive strain to investigate responses from the human cholangiocyte to *H. pylori*. However, repeat assays using other CagA positive strains such as P12 [83] could buttress the present findings and, as well, exclude strain-specific effects. Furthermore, inclusion of a CagA negative and/or *cag*PAI-dysfunctional strain such as SS1 would explore the specific contribution of CagA [84-86]. Second, induction of the hummingbird phenotype in gastric epithelial cells by the CagA+ *H. pylori* is dependent on the number and type of EPIYA phosphorylation site repeats [73, 87]. To confirm that Cag A+ *H. pylori* can also induce a hummingbird-like phenotype of the cholangiocyte, a demonstration that H69 cells can be infected by the *H. pylori*, and the *cag*PAI is functional during the infection, as established by detection of phosphorylated CagA, would be needed [88-90].

The present findings supported the hypothesis that opisthorchiasis and *H. pylori* together may hasten or even synergize the malignant transformation of cholangiocytes [2, 3, 91]. By contrast, co-infections of *Helicobacter* species including *H. pylori* and some other helminths have generally been associated with diminished risk of *H. pylori*-associated gastric carcinoma [92-94]. Notably, concurrent infection on mice with an intestinal nematode modulates inflammation, induces a Th2-polarizing cytokine phenotype with concomitant downmodulation of Th1 and the gastric immune responses, and reduces *Helicobacter*-induced gastric atrophy [95]. Nonetheless, given that CagA^+^ *H. pylori* stimulated EMT in cholangiocytes and which, in turn, also suggests a role in the underlying fibrosis [96-98] and metastasis [99-101] of cholangiocarcinoma, an explanation for why infection with the liver fluke induces CCA might now be somewhat clearer – involvement by *H. pylori* and its virulence factors [102, 103].

## Conclusions

Infection with CagA^+^ *H. pylori* induced epithelial to mesenchymal transition in human cholangiocytes. Further investigation of the relationship between *O. viverrini* and *H. pylori* within the infected biliary tract is warranted. Studies on tumorigenicity of the *H. pylori*-transformed H69 cells in immune-suppressed mice would likely be informative.

## Abbreviations

CCA: cholangiocarcinoma
EMT: epithelial to mesenchymal transition
CagA: cytotoxin-associated gene A
ZEB1: zinc finger E-box-binding homeobox 1
JAM1: junctional adhesion molecule 1
MMP7: matrix metalloprotease 7
CD24: cluster of differentiation marker 24
CD44: cluster of differentiation marker 44
MOI: multiplicity of infection
RTCA: real time cell analysis
CI: cell index
UC: upper chamber (of Boyden chamber)
LC: lower chamber

## Competing Financial Interests

The authors declare there were no competing financial interests.

## Acknowledgments

P.T. acknowledges support as a PhD research scholar from the Thailand Research Fund under the Royal Golden Jubilee scholars program (grant number PHD/0013/2555); B.S. acknowledges support from Thailand Research Fund Senior Research Scholar. This work was supported by the National Health Security Office, Thailand, the Higher Education Research Promotion and National Research University Project of Thailand, Office of the Higher Education Commission, through the Health Cluster (SHeP-GMS), the Faculty of Medicine, Khon Kaen University, Thailand (award number I56110), and the Thailand Research Fund under the TRF Senior Research Scholar (RTA 5680006); the National Research Council of Thailand. The National Institute of Allergy and Infectious Diseases (NIAID), Tropical Medicine Research Center award number P50AI098639, The National Cancer Institute (NCI), award number R01CA164719, and the United States Army Medical Research and Materiel Command (USAMRMC), contract number W81XWH-12-C-0267 also provided support. The content is solely the responsibility of the authors and does not necessarily represent the official views of the funders including USAMRMC, NIAID, NCI or the NIH. This manuscript has been released as a pre-print at *bioRxiv* [Thanaphongdecha et al [104]].

## Author contributions

P.T., S.S., C.P., Y.C., J.F., B.S. and P.J.B. conceived and designed the study. P.T., S.K., V.M., W.I. performed the experiments. P.T., W.I., B.S. and PJB analyzed and interpreted the findings. P.T., B.S., W.I., S.S., and PJB wrote the manuscript. All authors read and approved the final version of the paper.

## Supporting information

**Figure S1.**
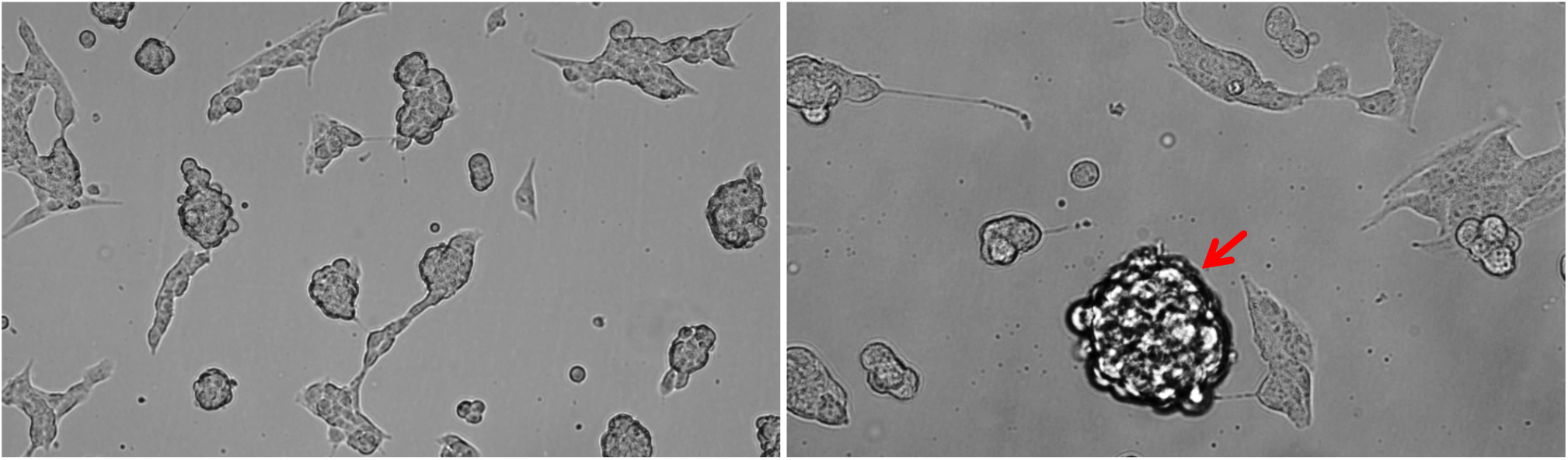
TGF-β promoted nodule formation. TGF-β, an EMT-stimulant, induced nodule formation of H69 cholangiocytes. Cells were cultured in the presence of 5 ng/ml TGF-β in serum- and hormone-free medium for 24 hours. Micrographs at 0 (left) and 24 (right) hours; 5×magnification.

**Figure S2.**
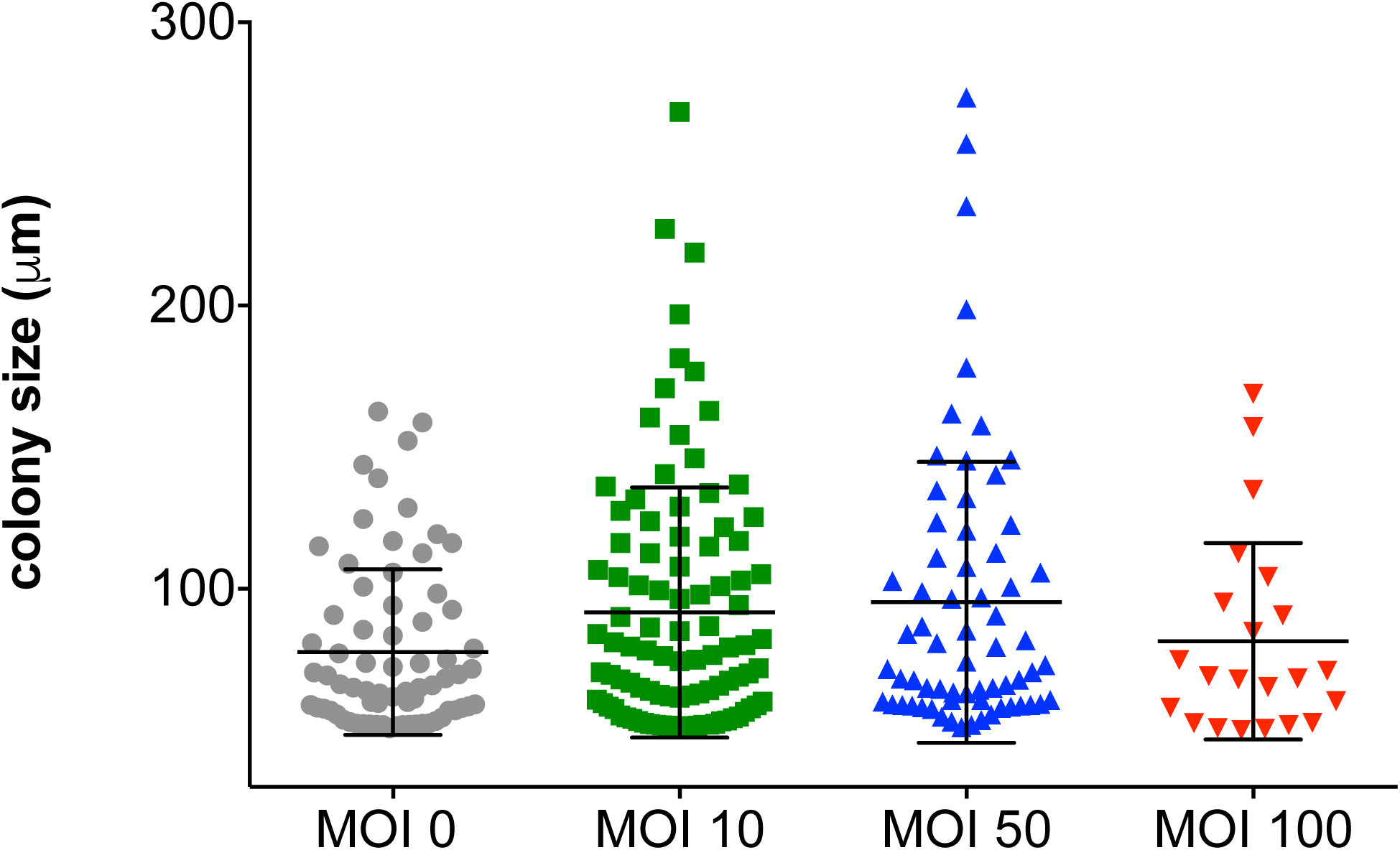
Size of colonies of cholangiocytes in soft agar. Colonies with a diameter ≥ 50 µm were counted and the diameter measured; mean values were 77.63, 91.62, 95.25 and 81.46 µm diameter at MOI of 0, 10, 50, and 100, respectively.

**Figure S3.**
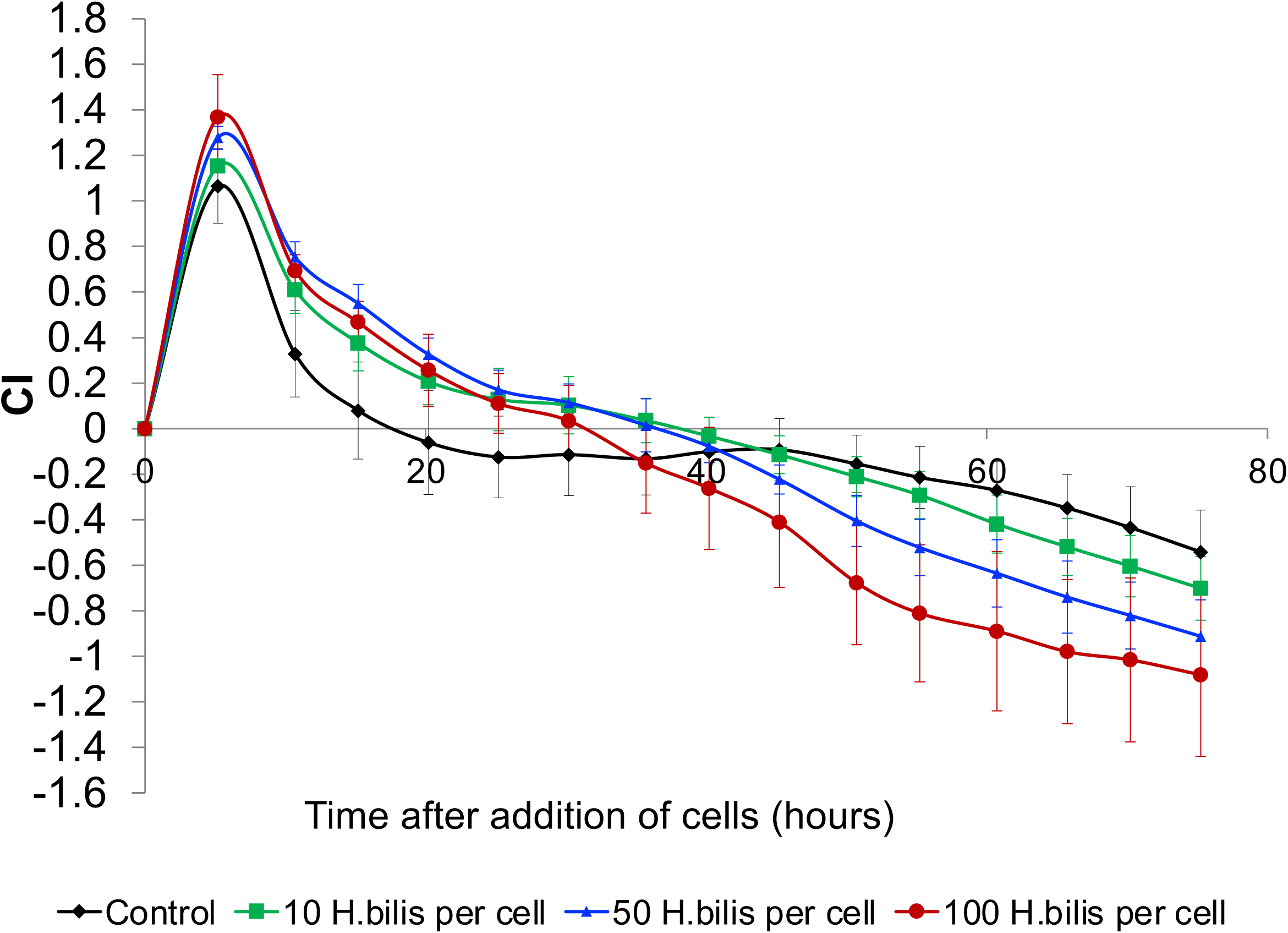
Growth of *Helicobacter bilis*-infected cholangiocytes as determined by the xCELLigence approach. H69 cells were exposed to increasing numbers of *H. bilis* (ATCC 43879) at MOI of 0, 10, 50, and 100. Cell growth was inhibited when compared to the uninfected cells, at all MOIs. Assay was performed in E-plates in minimal (serum-depleted) medium [36].

**Table S1.**
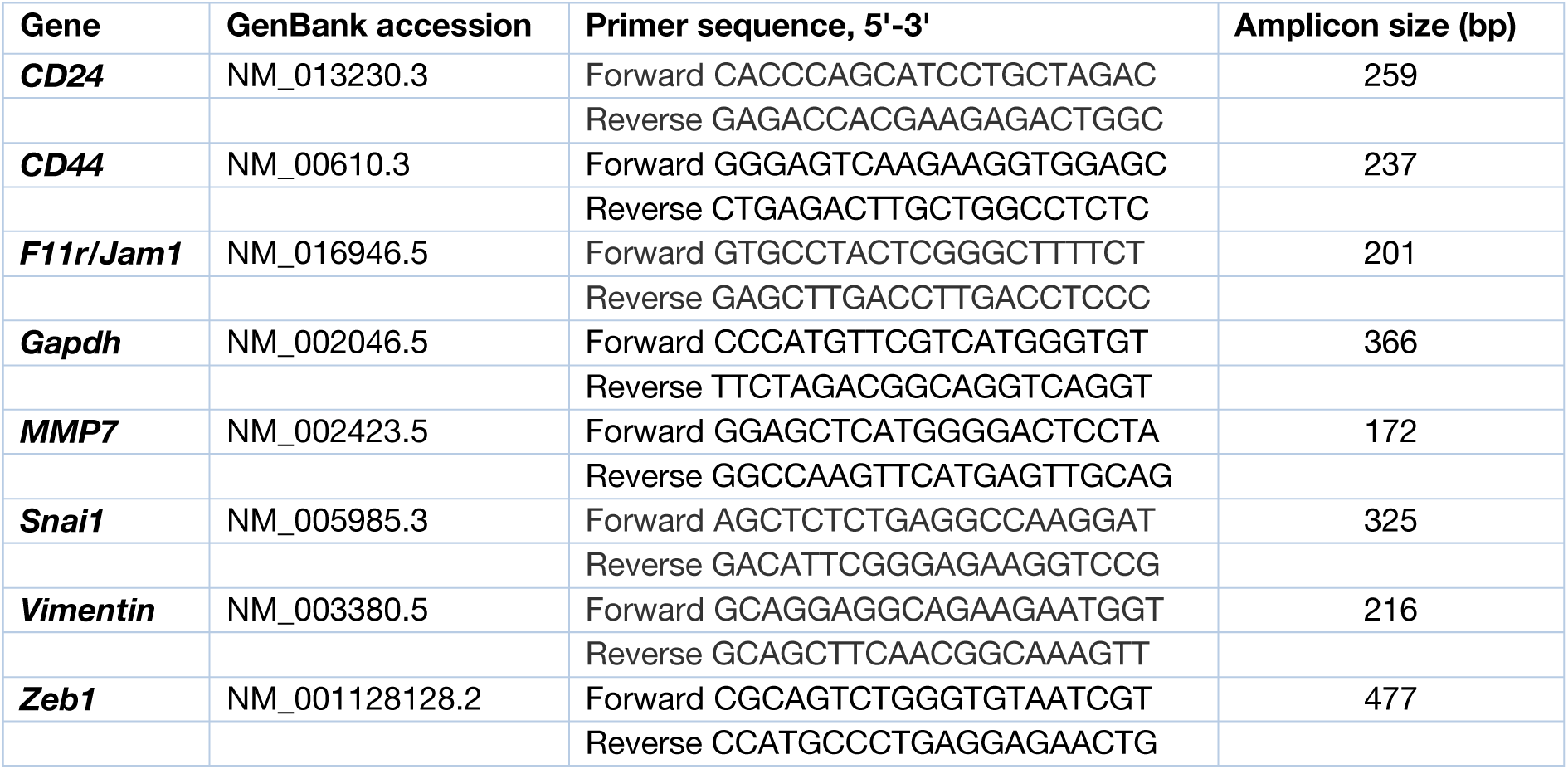
Nucleotide sequences of primers specific for human EMT-associated and cancer stem cell maker genes.

## Notes

### Competing Interest Statement

The authors have declared no competing interest.

### Summary of Updates

Updated results, figures and discussion

## References

1. Deenonpoe, R., et al., The carcinogenic liver fluke Opisthorchis viverrini is a reservoir for species of Helicobacter. Asian Pac J Cancer Prev, 2015. 16(5): p. 1751–8.

2. Deenonpoe, R., et al., Elevated prevalence of Helicobacter species and virulence factors in opisthorchiasis and associated hepatobiliary disease. Sci Rep, 2017. 7: p. 42744.

3. Sripa, B., R. Deenonpoe, and P.J. Brindley, Co-infections with liver fluke and Helicobacter species: A paradigm change in pathogenesis of opisthorchiasis and cholangiocarcinoma? Parasitol Int, 2017. 66(4): p. 383–389.

4. Boonyanugomol, W., et al., Effects of Helicobacter pylori gamma-glutamyltranspeptidase on apoptosis and inflammation in human biliary cells. Dig Dis Sci, 2012. 57(10): p. 2615–24.

5. Boonyanugomol, W., et al., Helicobacter pylori in Thai patients with cholangiocarcinoma and its association with biliary inflammation and proliferation. HPB (Oxford), 2012. 14(3): p. 177–84.

6. Humans, I.W.G.o.t.E.o.C.R.t., Biological agents. Volume 100 B. A review of human carcinogens. IARC Monogr Eval Carcinog Risks Hum, 2012. 100(Pt B): p. 1–441.

7. Sungkasubun, P., et al., Ultrasound screening for cholangiocarcinoma could detect premalignant lesions and early-stage diseases with survival benefits: a population-based prospective study of 4,225 subjects in an endemic area. BMC Cancer, 2016. 16: p. 346.

8. Khuntikeo, N., et al., A Comprehensive Public Health Conceptual Framework and Strategy to Effectively Combat Cholangiocarcinoma in Thailand. PLoS Negl Trop Dis, 2016. 10(1): p. e0004293.

9. Aye Soukhathammavong, P., et al., Subtle to severe hepatobiliary morbidity in Opisthorchis viverrini endemic settings in southern Laos. Acta Trop, 2015. 141(Pt B): p. 303–9.

10. Segura-Lopez, F.K., A. Guitron-Cantu, and J. Torres, Association between Helicobacter spp. infections and hepatobiliary malignancies: a review. World J Gastroenterol, 2015. 21(5): p. 1414–23.

11. Boonyanugomol, W., et al., Molecular analysis of Helicobacter pylori virulent-associated genes in hepatobiliary patients. HPB (Oxford), 2012. 14(11): p. 754–63.

12. Pisani, P., et al., Cross-reactivity between immune responses to Helicobacter bilis and Helicobacter pylori in a population in Thailand at high risk of developing cholangiocarcinoma. Clin Vaccine Immunol, 2008. 15(9): p. 1363–8.

13. Lvova, M.N., et al., Comparative histopathology of Opisthorchis felineus and Opisthorchis viverrini in a hamster model: an implication of high pathogenicity of the European liver fluke. Parasitol Int, 2012. 61(1): p. 167–72.

14. Mairiang, E., et al., Ultrasonography assessment of hepatobiliary abnormalities in 3359 subjects with Opisthorchis viverrini infection in endemic areas of Thailand. Parasitol Int, 2012. 61(1): p. 208–11.

15. Kao, C.Y., B.S. Sheu, and J.J. Wu, Helicobacter pylori infection: An overview of bacterial virulence factors and pathogenesis. Biomed J, 2016. 39(1): p. 14–23.

16. Cid, T.P., et al., Pathogenesis of Helicobacter pylori infection. Helicobacter, 2013. 18 Suppl 1: p. 12–7.

17. Hatakeyama, M., Helicobacter pylori CagA and gastric cancer: a paradigm for hit-and-run carcinogenesis. Cell Host Microbe, 2014. 15(3): p. 306–16.

18. Cover, T.L., D.B. Lacy, and M.D. Ohi, The Helicobacter pylori Cag Type IV Secretion System. Trends Microbiol, 2020. 28(8): p. 682–695.

19. Fox, J.G., et al., Hepatic Helicobacter species identified in bile and gallbladder tissue from Chileans with chronic cholecystitis. Gastroenterology, 1998. 114(4): p. 755–63.

20. Pellicano, R., et al., Helicobacter species sequences in liver samples from patients with and without hepatocellular carcinoma. World J Gastroenterol, 2004. 10(4): p. 598–601.

21. Nilsson, H.O., et al., Helicobacter species identified in liver from patients with cholangiocarcinoma and hepatocellular carcinoma. Gastroenterology, 2001. 120(1): p. 323–4.

22. Huang, Y., et al., Identification of helicobacter species in human liver samples from patients with primary hepatocellular carcinoma. J Clin Pathol, 2004. 57(12): p. 1273–7.

23. Knorr, J., et al., Classification of Helicobacter pylori Virulence Factors: Is CagA a Toxin or Not? Trends Microbiol, 2019. 27(9): p. 731–738.

24. Boonyanugomol, W., et al., Role of cagA-positive Helicobacter pylori on cell proliferation, apoptosis, and inflammation in biliary cells. Dig Dis Sci, 2011. 56(6): p. 1682–92.

25. Plieskatt, J.L., et al., Infection with the carcinogenic liver fluke Opisthorchis viverrini modifies intestinal and biliary microbiome. FASEB J, 2013. 27(11): p. 4572–84.

26. Boonyanugomol, W., et al., Helicobacter pylori cag pathogenicity island (cagPAI) involved in bacterial internalization and IL-8 induced responses via NOD1-and MyD88-dependent mechanisms in human biliary epithelial cells. PLoS One, 2013. 8(10): p. e77358.

27. Fung, C., et al., High-resolution mapping reveals that microniches in the gastric glands control Helicobacter pylori colonization of the stomach. PLoS Biol, 2019. 17(5): p. e3000231.

28. Lamouille, S., J. Xu, and R. Derynck, Molecular mechanisms of epithelial-mesenchymal transition. Nat Rev Mol Cell Biol, 2014. 15(3): p. 178–96.

29. Hatakeyama, M., Oncogenic mechanisms of the Helicobacter pylori CagA protein. Nat Rev Cancer, 2004. 4(9): p. 688–94.

30. Fox, J.G., Helicobacter bilis: bacterial provocateur orchestrates host immune responses to commensal flora in a model of inflammatory bowel disease. Gut, 2007. 56(7): p. 898–900.

31. Grubman, S.A., et al., Regulation of intracellular pH by immortalized human intrahepatic biliary epithelial cell lines. Am J Physiol, 1994. 266(6 Pt 1): p. G1060–70.

32. Park, J., G.J. Gores, and T. Patel, Lipopolysaccharide induces cholangiocyte proliferation via an interleukin-6-mediated activation of p44/p42 mitogen-activated protein kinase. Hepatology, 1999. 29(4): p. 1037–43.

33. Bairoch, A., The Cellosaurus, a Cell-Line Knowledge Resource. J Biomol Tech, 2018. 29(2): p. 25–38.

34. Shimizu, Y., et al., Two new human cholangiocarcinoma cell lines and their cytogenetics and responses to growth factors, hormones, cytokines or immunologic effector cells. Int J Cancer, 1992. 52(2): p. 252–60.

35. Han, C., et al., PPARgamma ligands inhibit cholangiocarcinoma cell growth through p53-dependent GADD45 and p21 pathway. Hepatology, 2003. 38(1): p. 167–77.

36. Smout, M.J., et al., Carcinogenic Parasite Secretes Growth Factor That Accelerates Wound Healing and Potentially Promotes Neoplasia. PLoS Pathog, 2015. 11(10): p. e1005209.

37. Arunsan, P., et al., Programmed knockout mutation of liver fluke granulin attenuates virulence of infection-induced hepatobiliary morbidity. Elife, 2019. 8.

38. Marshall, B.J. and J.R. Warren, Unidentified curved bacilli in the stomach of patients with gastritis and peptic ulceration. Lancet, 1984. 1(8390): p. 1311–5.

39. Dewhirst, F.E., et al., ‘Flexispira rappini’ strains represent at least 10 Helicobacter taxa. Int J Syst Evol Microbiol, 2000. 50 Pt 5: p. 1781–1787.

40. Fox, J.G., et al., Helicobacter bilis sp. nov., a novel Helicobacter species isolated from bile, livers, and intestines of aged, inbred mice. J Clin Microbiol, 1995. 33(2): p. 445–54.

41. Hachem, C.Y., et al., Comparison of agar based media for primary isolation of Helicobacter pylori. J Clin Pathol, 1995. 48(8): p. 714–6.

42. Jung, S.W., et al., Mechanism of antibacterial activity of liposomal linolenic acid against Helicobacter pylori. PLoS One, 2015. 10(3): p. e0116519.

43. Lin, S.N., et al., Helicobacter pylori heat-shock protein 60 induces production of the pro-inflammatory cytokine IL8 in monocytic cells. J Med Microbiol, 2005. 54(Pt 3): p. 225–233.

44. Pan, H., et al., A comparison of conventional methods for the quantification of bacterial cells after exposure to metal oxide nanoparticles. BMC Microbiol, 2014. 14: p. 222.

45. Masuda, M., et al., Tumor suppressor in lung cancer (TSLC)1 suppresses epithelial cell scattering and tubulogenesis. J Biol Chem, 2005. 280(51): p. 42164–71.

46. Bourzac, K.M., L.A. Satkamp, and K. Guillemin, The Helicobacter pylori cag pathogenicity island protein CagN is a bacterial membrane-associated protein that is processed at its C terminus. Infect Immun, 2006. 74(5): p. 2537–43.

47. Nagy, T.A., et al., Helicobacter pylori regulates cellular migration and apoptosis by activation of phosphatidylinositol 3-kinase signaling. J Infect Dis, 2009. 199(5): p. 641–51.

48. Bindschadler, M. and J.L. McGrath, Sheet migration by wounded monolayers as an emergent property of single-cell dynamics. J Cell Sci, 2007. 120(Pt 5): p. 876–84.

49. Liang, C.C., A.Y. Park, and J.L. Guan, In vitro scratch assay: a convenient and inexpensive method for analysis of cell migration in vitro. Nat Protoc, 2007. 2(2): p. 329–33.

50. Livak, K.J. and T.D. Schmittgen, Analysis of relative gene expression data using real-time quantitative PCR and the 2(-Delta Delta C(T)) Method. Methods, 2001. 25(4): p. 402–8.

51. Ye, J., et al., Primer-BLAST: a tool to design target-specific primers for polymerase chain reaction. BMC Bioinformatics, 2012. 13: p. 134.

52. Arunsan, P., et al., Liver fluke granulin promotes extracellular vesicle-mediated crosstalk and cellular microenvironment conducive to cholangiocarcinoma. Neoplasia, 2020. 22(5): p. 203–216.

53. Ke, N., et al., The xCELLigence system for real-time and label-free monitoring of cell viability. Methods Mol Biol, 2011. 740: p. 33–43.

54. Dowling, C.M., C. Herranz Ors, and P.A. Kiely, Using real-time impedance-based assays to monitor the effects of fibroblast-derived media on the adhesion, proliferation, migration and invasion of colon cancer cells. Biosci Rep, 2014. 34(4).

55. Matchimakul, P., et al., Apoptosis of cholangiocytes modulated by thioredoxin of carcinogenic liver fluke. Int J Biochem Cell Biol, 2015. 65: p. 72–80.

56. Rinaldi, G., et al., Cytometric analysis, genetic manipulation and antibiotic selection of the snail embryonic cell line Bge from Biomphalaria glabrata, the intermediate host of Schistosoma mansoni. Int J Parasitol, 2015. 45(8): p. 527–35.

57. Solly, K., et al., Application of real-time cell electronic sensing (RT-CES) technology to cell-based assays. Assay Drug Dev Technol, 2004. 2(4): p. 363–72.

58. Xie, H., et al., Novel functions and targets of miR-944 in human cervical cancer cells. Int J Cancer, 2015. 136(5): p. E230–41.

59. Horibata, S., et al., Utilization of the Soft Agar Colony Formation Assay to Identify Inhibitors of Tumorigenicity in Breast Cancer Cells. J Vis Exp, 2015(99): p. e52727.

60. Zeisberg, M. and E.G. Neilson, Biomarkers for epithelial-mesenchymal transitions. J Clin Invest, 2009. 119(6): p. 1429–37.

61. Nieto, M.A., The snail superfamily of zinc-finger transcription factors. Nat Rev Mol Cell Biol, 2002. 3(3): p. 155–66.

62. Boulay, J.L., C. Dennefeld, and A. Alberga, The Drosophila developmental gene snail encodes a protein with nucleic acid binding fingers. Nature, 1987. 330(6146): p. 395–8.

63. Nguyen, P.H., et al., Characterization of Biomarkers of Tumorigenic and Chemoresistant Cancer Stem Cells in Human Gastric Carcinoma. Clin Cancer Res, 2017. 23(6): p. 1586–1597.

64. Shomer, N.H., et al., Helicobacter bilis-induced inflammatory bowel disease in scid mice with defined flora. Infect Immun, 1997. 65(11): p. 4858–64.

65. Flahou, B., et al., The Other Helicobacters. Helicobacter, 2015. 20 Suppl 1: p. 62–7.

66. Cover, T.L., Helicobacter pylori Diversity and Gastric Cancer Risk. MBio, 2016. 7(1): p. e01869–15.

67. Marshall, B.J., The pathogenesis of non-ulcer dyspepsia. Med J Aust, 1985. 143(7): p. 319.

68. Gaynor, E.C. and C.M. Szymanski, The 30th anniversary of Campylobacter, Helicobacter, and Related Organisms workshops-what have we learned in three decades? Front Cell Infect Microbiol, 2012. 2: p. 20.

69. Sheh, A. and J.G. Fox, The role of the gastrointestinal microbiome in Helicobacter pylori pathogenesis. Gut Microbes, 2013. 4(6): p. 505–31.

70. Kienesberger, S., et al., Gastric Helicobacter pylori Infection Affects Local and Distant Microbial Populations and Host Responses. Cell Rep, 2016. 14(6): p. 1395–407.

71. Cover, T.L. and M.J. Blaser, Helicobacter pylori in health and disease. Gastroenterology, 2009. 136(6): p. 1863–73.

72. Saadat, I., et al., Helicobacter pylori CagA targets PAR1/MARK kinase to disrupt epithelial cell polarity. Nature, 2007. 447(7142): p. 330–3.

73. Hatakeyama, M. and H. Higashi, Helicobacter pylori CagA: a new paradigm for bacterial carcinogenesis. Cancer Sci, 2005. 96(12): p. 835–43.

74. Yamaoka, Y., Mechanisms of disease: Helicobacter pylori virulence factors. Nat Rev Gastroenterol Hepatol, 2010. 7(11): p. 629–41.

75. Murphy, G., et al., Association of seropositivity to Helicobacter species and biliary tract cancer in the ATBC study. Hepatology, 2014. 60(6): p. 1963–71.

76. Mateos-Munoz, B., et al., Enterohepatic Helicobacter other than Helicobacter pylori. Rev Esp Enferm Dig, 2013. 105(8): p. 477–84.

77. Zhou, D., et al., Infections of Helicobacter spp. in the biliary system are associated with biliary tract cancer: a meta-analysis. Eur J Gastroenterol Hepatol, 2013. 25(4): p. 447–54.

78. Pellicano, R., et al., Helicobacter species and liver diseases: association or causation? Lancet Infect Dis, 2008. 8(4): p. 254–60.

79. Wroblewski, L.E., R.M. Peek, Jr., and K.T. Wilson, Helicobacter pylori and gastric cancer: factors that modulate disease risk. Clin Microbiol Rev, 2010. 23(4): p. 713–39.

80. Segal, E.D., et al., Altered states: involvement of phosphorylated CagA in the induction of host cellular growth changes by Helicobacter pylori. Proc Natl Acad Sci U S A, 1999. 96(25): p. 14559–64.

81. Klonisch, T., et al., Cancer stem cell markers in common cancers - therapeutic implications. Trends Mol Med, 2008. 14(10): p. 450–60.

82. Simpson, C.D., K. Anyiwe, and A.D. Schimmer, Anoikis resistance and tumor metastasis. Cancer Lett, 2008. 272(2): p. 177–85.

83. Fischer, W., et al., Strain-specific genes of Helicobacter pylori: genome evolution driven by a novel type IV secretion system and genomic island transfer. Nucleic Acids Res, 2010. 38(18): p. 6089–101.

84. Lee, A., et al., A standardized mouse model of Helicobacter pylori infection: introducing the Sydney strain. Gastroenterology, 1997. 112(4): p. 1386–97.

85. Nolan, K.J., et al., In vivo behavior of a Helicobacter pylori SS1 nixA mutant with reduced urease activity. Infect Immun, 2002. 70(2): p. 685–91.

86. Kundu, P., et al., Cag pathogenicity island-independent up-regulation of matrix metalloproteinases-9 and -2 secretion and expression in mice by Helicobacter pylori infection. J Biol Chem, 2006. 281(45): p. 34651–62.

87. Higashi, H., et al., EPIYA motif is a membrane-targeting signal of Helicobacter pylori virulence factor CagA in mammalian cells. J Biol Chem, 2005. 280(24): p. 23130–7.

88. Ren, S., et al., Structural basis and functional consequence of Helicobacter pylori CagA multimerization in cells. J Biol Chem, 2006. 281(43): p. 32344–52.

89. Backert, S., N. Tegtmeyer, and W. Fischer, Composition, structure and function of the Helicobacter pylori cag pathogenicity island encoded type IV secretion system. Future Microbiol, 2015. 10(6): p. 955–65.

90. Tohidpour, A., CagA-mediated pathogenesis of Helicobacter pylori. Microb Pathog, 2016. 93: p. 44–55.

91. Itthitaetrakool, U., et al., Chronic Opisthorchis viverrini Infection Changes the Liver Microbiome and Promotes Helicobacter Growth. PLoS One, 2016. 11(11): p. e0165798.

92. Holcombe, C., Helicobacter pylori: the African enigma. Gut, 1992. 33(4): p. 429–31.

93. Du, Y., et al., Helicobacter pylori and Schistosoma japonicum co-infection in a Chinese population: helminth infection alters humoral responses to H. pylori and serum pepsinogen I/II ratio. Microbes Infect, 2006. 8(1): p. 52–60.

94. Whary, M.T., et al., Intestinal helminthiasis in Colombian children promotes a Th2 response to Helicobacter pylori: possible implications for gastric carcinogenesis. Cancer Epidemiol Biomarkers Prev, 2005. 14(6): p. 1464–9.

95. Fox, J.G., et al., Concurrent enteric helminth infection modulates inflammation and gastric immune responses and reduces helicobacter-induced gastric atrophy. Nat Med, 2000. 6(5): p. 536–42.

96. Fragiadaki, M. and R.M. Mason, Epithelial-mesenchymal transition in renal fibrosis - evidence for and against. Int J Exp Pathol, 2011. 92(3): p. 143–50.

97. Rout-Pitt, N., et al., Epithelial mesenchymal transition (EMT): a universal process in lung diseases with implications for cystic fibrosis pathophysiology. Respir Res, 2018. 19(1): p. 136.

98. Stone, R.C., et al., Epithelial-mesenchymal transition in tissue repair and fibrosis. Cell Tissue Res, 2016. 365(3): p. 495–506.

99. Kalluri, R. and R.A. Weinberg, The basics of epithelial-mesenchymal transition. J Clin Invest, 2009. 119(6): p. 1420–8.

100. Li, L. and W. Li, Epithelial-mesenchymal transition in human cancer: comprehensive reprogramming of metabolism, epigenetics, and differentiation. Pharmacol Ther, 2015. 150: p. 33–46.

101. Sciacovelli, M. and C. Frezza, Metabolic reprogramming and epithelial-to-mesenchymal transition in cancer. FEBS J, 2017. 284(19): p. 3132–3144.

102. Brindley, P.J., J.M. da Costa, and B. Sripa, Why does infection with some helminths cause cancer? Trends Cancer, 2015. 1(3): p. 174–182.

103. Brindley, P.J. and A. Loukas, Helminth infection-induced malignancy. PLoS Pathog, 2017. 13(7): p. e1006393.

104. Thanaphongdecha, P.K. S.E., Ittiprasrt, W., Mann, V.H., Chamgramol, Y., Pairojkul, C., Fox, J.G., Suttiprapa, S., Sripa, B., Brindley, P.J., Infection with CagA+ Helicobacter pylori induces epithelial to mesenchymal transition in human cholangiocytes. bioRxiv, 2020.

